# Analyzing and interpreting DNA double-strand break sequencing data

**DOI:** 10.1101/2020.03.05.977801

**Authors:** Abhishek Mitra, Norbert Dojer, Bernard Fongang, Jules Nde, Yingjie Zhu, Maga Rowicka

**Affiliations:** Department of Biochemistry and Molecular Biology, University of Texas Medical Branch at Galveston, Galveston, USA; Institute of Informatics, University of Warsaw, Warsaw, Poland; Institute for Translational Sciences, University of Texas Medical Branch at Galveston, Galveston, Texas, USA; Sealy Center for Molecular Medicine, University of Texas Medical Branch at Galveston, Galveston, Texas, USA; Sealy Center for Structural Biology and Molecular Biophysics, University of Texas Medical Branch at Galveston, Galveston, Texas, USA

## Abstract

DNA double-strand breaks (DSBs), are a major threat to genomic stability and may lead to cancer. Several technologies to accurately detect DSBs genome-wide have been developed recently, but still lacking publicly available tools for analysis of the resulting data. Here, we present a step-by-step iSeq package (http://breakome.utmb.edu/software.html), custom designed for analysis and interpretation of DSB-sequencing data. iSeq performs barcode trimming and read counting, and identifies DSB-enriched regions by statistical test and annotate them to the desired genomic features. Applying this package, users can identify and annotate DSB-enriched regions from base pair (eg. Cas9 cleavage sites) up to megabase (eg. DNA replication stress-induced) resolution, and if possible quantify DSB frequencies per cell genome-wide by combining with qDSB-Seq. iSeq can be used for any sequencing-based DSB detection techniques. The analysis for Steps 1-19 can be performed within ~4 hours.

## Introduction

DNA double-strand breaks (DSBs) are the most genotoxic form of DNA damage. There is extensive evidence that DSBs are a major threat to genomic stability, responsible for chromosomal instability and rearrangements, and arise through the direct action of ionizing radiation or chemicals (including many chemotherapy drugs)^1^, as well as during replication^2^. A better understanding of their formation and occurrence is paramount to effective prevention and treatment of the associated diseases, including cancer^3^. Despite this importance, knowledge of how the genome breaks in response to various stressors is still sorely lacking. This deficit has been a direct result of the absence of critical research tools. Recently, several technologies were developed to accurately detect DSBs throughout the genome by sequencing^4–8^, however, there is still lacking advanced statistical methods and tools for analysis and interpretation of these data. Here we present the novel advanced statistical methods implemented as user-friendly software tools that overcome this final challenge.

We developed a method, BLESS, to label DSBs *in situ* followed by deep sequencing in 2013^4^, which allows the study of DSBs with nucleotide resolution^9–11^. Several modifications of our method, such as i-BLESS, END-Seq, Break-Seq or DSB-capture have been published^5–8^. Working with more than 100 samples from DSB sequencing using these various methods^4–11^, we developed the first dedicated software suite for analysis of DSB sequencing data, especially customized for this unusual, emerging data type. The paucity of DSBs makes their analysis particularly challenging and results in weak signal. Therefore, normalization is crucial. Moreover, DSB sequencing leads to different patterns than, for example ChIP-seq, so using software created for ChIP-seq analysis, such as MACS^12^, is inappropriate for some DSB patterns, for example, replication-related DSBs. Moreover, DSB sequencing detects the end of DNA fragment rather than its body, so the coverage cannot be regularly counted by common software. Some standard practices such as removing duplicate reads, often leads to incorrect results in DSB sequencing. Therefore, we developed a dedicated pipeline, Instant Sequencing (iSeq), for DSB sequencing data analysis that starts from pre-processing the raw sequencing reads and ends with high-level data interpretation. For instance, our software is capable of calculating enrichment of detected DSBs in available annotated files, and rigorously estimating the chances for any given gene to have DSBs within its body or its promoter under any given conditions. Our approach is also unique in that it provides multiscale analysis, starting from base pairs resolution and gradually progressing up to mega base resolution, and automatically generates reports about trends across scales. In addition, we introduce the calculation of DSBs per cell and across-samples comparison based on qDSB-Seq data and software^13^. The software of iSeq and qDSB-Seq can be find on the links of http://breakome.utmb.edu/software.html.

## Applications

Since publication of our DSB detection method^4^, we originally applied the suggested methods to study genomic landscape of sensitivity to replication stress by identifying aphidicolin-sensitive regions (ARSs). To evaluate the impact of replication stress to genome instability, we exposed HeLa cells to aphidicolin, which is a DNA polymerase inhibitor and can induce replication-fork stalling without arresting progression in S-phase. We detected significant accumulation of breaks and increased the amount of labeled DSBs captured by BLESS^4^. We identified ARSs by calculating enrichment *P* values based on the hypergeometric distribution and their corrections based on the Benjamini-Hochberg approach for multiple hypothesis testing. We noticed that ARSs were not uniformly distributed on the chromosomes, so we characterized the distribution of ARSs by genomic annotation. Further, we found that these ARSs overrepresented in genes and enriched in satellite repeats, suggesting that the genomic instability from replication stress could be related with the transcription-replication collision and the special structure of repeats.

In another application of BLESS, we reported the roles of Ssb1 and Ssb2 proteins in the regulation of the DNA damage response^11^. By application enrichment analysis in iSeq, we found that cells lacking *Ssb1/Ssb2* induce genome-wide double-strand breaks enriched at CpG islands and transfer RNAs, especially for those located on Ssb1 and Ssb2 binding sites. Other applications include investigating Homologous Recombination (HR)-prone DSBs and Non-homologous End Joining (NHEJ)-prone DSBs^10^, studying the on-target and off-target effects of the Cas9 endonuclease^14^.

Recently, we presented a general method, qDSB-Seq^13^, to quantify DSBs genome-wide, which provides both DSB frequencies per cell and their precise genomic coordinates. qDSB-Seq has been applied to BLESS^4^ and i-BLESS^5^, it is also available for END-Seq^6^, Break-Seq^7^, DSBCapture^8^, BLISS^15^, and so on. In this work, we induced spike-in DSBs by a site-specific endonuclease and used these induced DSBs to quantify studied DSBs. qDSB-Seq method has been validated in more than 35 experiments, including yeast cells in untreated G1-phase, Zeocin-treated G1-phase, *pif1-m2* mutant G1-phase, hydroxyurea (HU)-treated S-phase, release from HU, camptothecin-treated S-phase, and also human DIvA cells upon 4-hydroxytamoxifen (4OHT) treatment. In these samples, qDSB-Seq software we developed was used for calculating the total number of DSBs per cell. It also computes DSBs at a specific locus or a region, for example, replication fork barrier or replication fork origin. In combination with hypergeometric test in iSeq package, qDSB-Seq software quantified 1-ended DSBs induced from DNA replication stress.

In this protocol, we describe and discuss in detail how the computational framework can be better utilized to identify DSB-enriched regions and where they are prone to locate on the genome. We also present how to identify DSBs from enzyme cutting sites, including off-target sites, how to quantify DSBs based on endonuclease-digested DNA. Although the example protocol described was on the basis of BLESS^4^ and i-BLESS^5^, it can be also used for other DSB sequencing methods with minor changes in barcode trimming stage.

## Overview of the procedure

### Data processing and normalization

Data processing examines sequencing quality and trim barcode sequences from DSB sequencing reads (**Fig. 1**). The processed reads are further mapped to reference genome and used for reads counting. Frequently used tools, for example, SAMtools^16^ and bedtools (https://bedtools.readthedocs.io/en/latest/) count sequencing coverage from every nucleotides of a sequencing read, however, the read coverage of DSB sequencing should be counted from the first nucleotide of a sequencing read. Our tool provides read counts from the first nucleotide of a sequencing read and both Watson and Crick strands. The next step is comprehensive data normalization by either Copy Number Variation (CNV) or DSB sequencing data of other samples. Since DSBs are very rare events, the resulting sequencing signal is often weaker than experimental biases, and therefore proper normalization is crucial. For example, without such normalization, in cancer cell lines, breaks may be discovered predominantly in amplified regions (**Supplementary Fig. 1**). Some common treatments (e.g. aphidicolin) are known to affect replication speed, which biases sequencing results in a manner similar to CNV. When the appropriate controls are also sequenced, the software corrects for CNV, treatment effects and other artifacts of sequencing and sample preparation (chromatin accessibility, PCR-bias, etc.), for every “treatment (T) vs. control (C)” experiment.

**Fig. 1.**
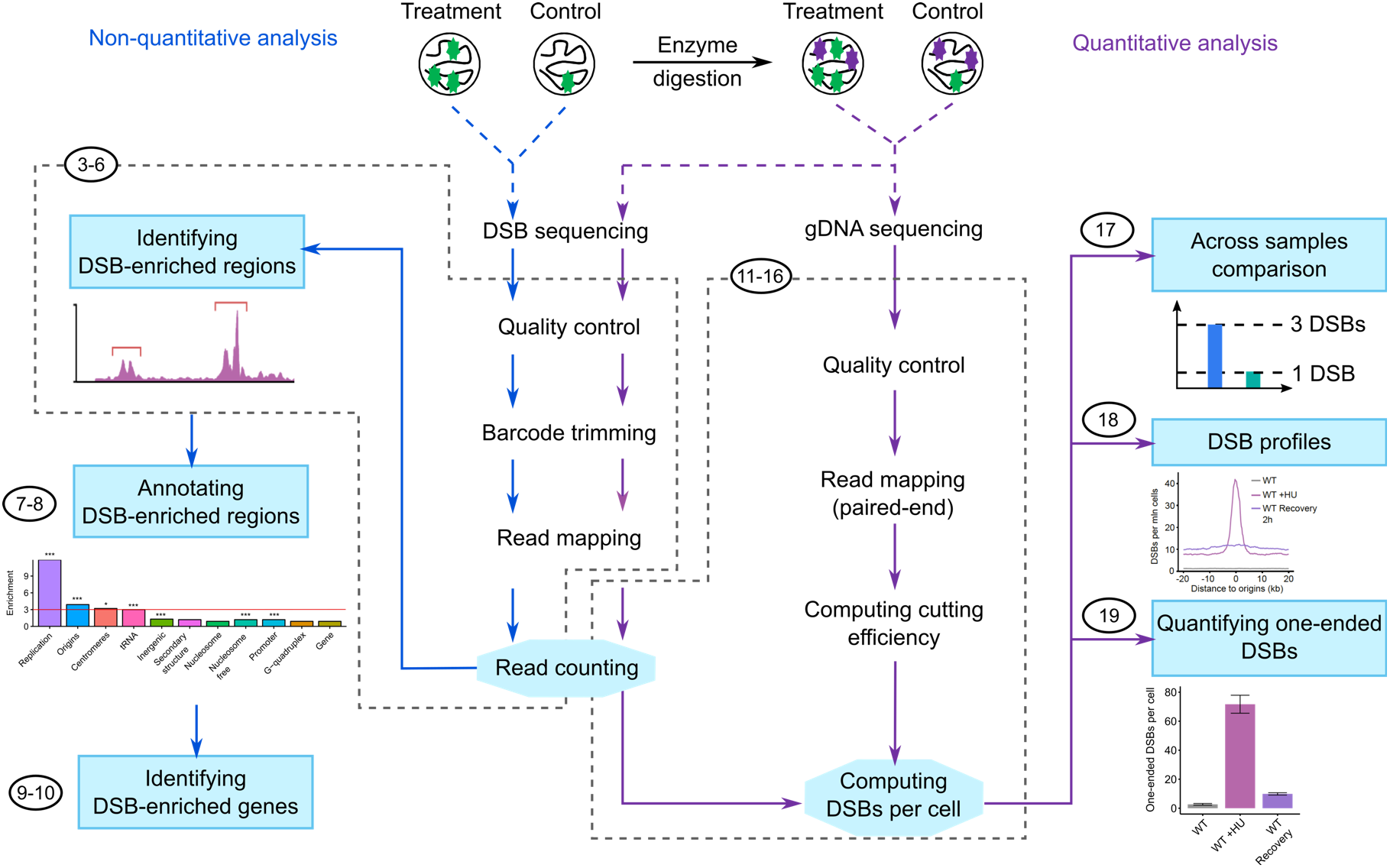
Overview of Instant Sequencing (iSeq) data interpretation and analysis pipeline. The number of steps is shown.

### Identification of DSB-enriched regions

To identify genomic regions where are DSB-enriched, we developed a statistical method to find regions with significant difference between treatment and control samples. The genome was divided into continuous genomic windows with a constant number of mappable nucleotides. The number of sequencing reads in the window was counted for the statistical analysis. We used hypergeometric distribution to estimate *P* values, which were further corrected with multiple hypothesis-testing corrections (Benjamin-Hochberg or Bonferroni approaches). By this way, we account for the variation in mappability along the genome (and thus the expected number of sequencing reads), and produce more statistically robust comparisons of the number of reads between different windows. We verified that such windows are not biased toward detecting particular genomic features^4^.

We have shown^4^ that except for chromosome-wide DSB frequency, which is fairly constant, DSB-prone regions are distributed very unevenly in the human genome (**Supplementary Fig. 2**). At higher resolution (top panel of **Supplementary Fig. 2**) DSB-prone regions can be found both in highly breakable and non-breakable lower resolution regions (pink and teal regions in the middle panel of **Supplementary Fig. 2**). Moreover, low- and high-resolution DSB-prone regions exhibit very different properties, as shown in **Fig. 2**, therefore multi-scale analysis is very important. Another reason to perform multi-scale analysis is to eliminate spurious enrichments, resulting from a specific choice of parameters. In multi-scale analysis, they will be easy to detect, they would manifest themselves and an enrichments or depletions observable at only one specific resolution, but not at any similar ones, unlike genuine enrichments, that would vary smoothly between resolutions (**Fig. 2**).

**Fig. 2.**
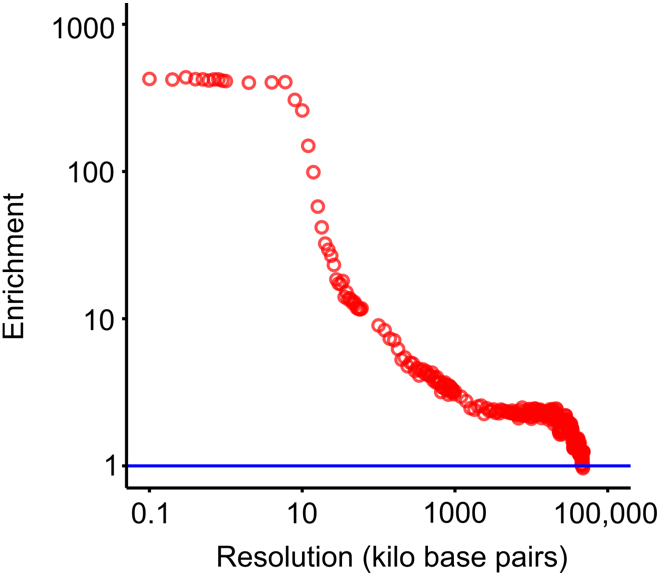
Log-log plot of dependence of alpha satellite repeats enrichment of DSBPRs (vertical axis) on the resolution of DSB detection (horizontal axis). Blue horizontal line denotes no enrichment. Note almost 1000-fold change in enrichment across resolutions.

**Fig. 3.**
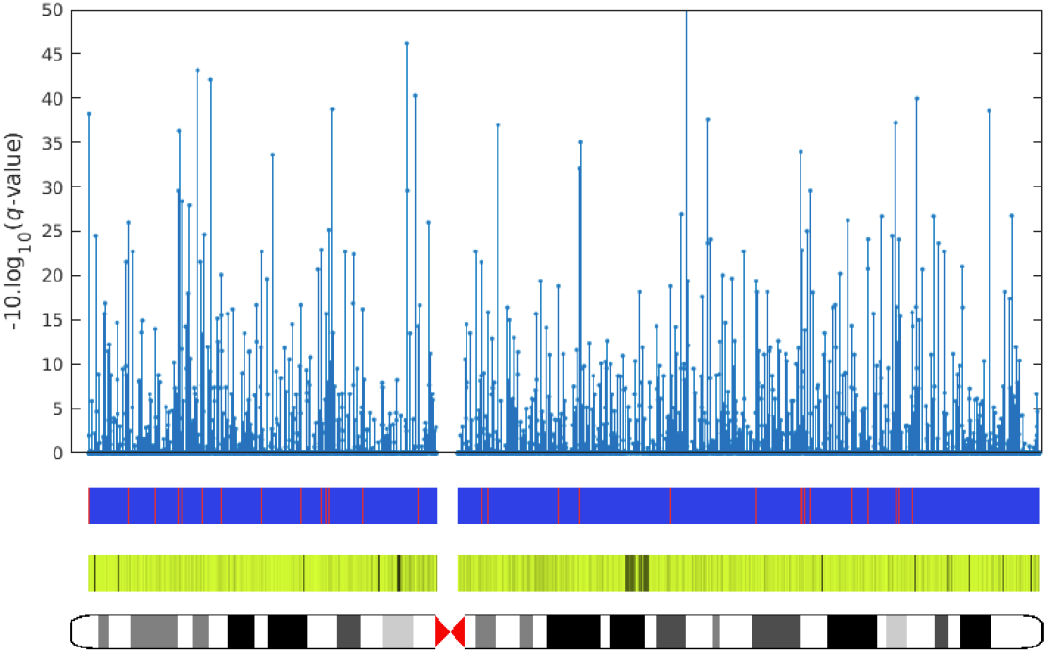
Sample plot of multiscale fragility maps generated by iSeq in Step 6. Intervals on chromosome X in aphidicolin-HeLa cells are shown. From bottom: 1) chromosome ideogram (centromere is denoted by red hourglass); 2) mapability (black denotes 0%); 3) breaking probability (red denotes 100%, blue 0%) for 48 kilobase resolution; 4) −10Log_10_(q-value) for DSB presence at 48 kilobase resolution (blue stem plot).

To identify DSB-enriched genes, we used similar method in identifying DSB-enriched regions. Differently, the number of reads in mappable gene regions rather than genomic windows was counted for the hypergeometric test. In the same manner, the *P* values for each gene were corrected by Benjamin-Hochberg correction. Thus, we can find genes with significant DNA damage in different conditions.

### Annotation of DSB-enriched regions

To annotate DSB-enriched regions to given genomic features, we developed a feature score to evaluate the relationship between DSB-enriched regions and features. First, we computed observed overlap, which is the proportion of mappable nucleotides belonging to both DSB-enriched regions and given genomic regions, and expected overlap, which is the proportion of mappable nucleotides belonging to both genomic regions and the given feature. The feature score is the proportion between observed overlap and expected overlap. We used this score to evaluate DSB-enriched regions are simultaneously enriched in the given feature (feature score > 1) or depleted (feature score < 1). Second, to estimate *P* value for the feature enrichment or depletion, we used permutations of DSB-enriched regions among the windows considered and utilized them to calculate the empirical distribution of the annotation score under the null hypothesis that the given feature and DSB-enriched regions are independently distributed in the genome.

### DSB quantification

We developed qDSB-Seq^13^ and its computational solution to quantify DSBs per cell by inducing measurable DSBs by a site-specific endonuclease. The assumption of qDSB-Seq quantification is that the number of DSB sequencing reads is proportional to the underlying DSB frequency. When DSBs at pre-determined genomic loci are induced and measured, we can find the relationship between DSB sequencing reads and DSB frequencies. In the qDSB-Seq approach, we induced spike-in DSBs by a restriction endonuclease. Then, their DSB frequencies are determined from genomic DNA (gDNA) sequencing data, or from quantitative PCR data. The number of DSBs at cutting sites is computed from DSB frequencies by multiplying the number of cutting sites. Finally, the average number of studied DSBs is estimated based on the proportion measured from restriction enzyme digestion. To eliminate background of the estimated DSB number, which could be caused by genome fragmentation in gDNA sequencing, we estimated the background level of DSBs from gDNA data by randomly selecting genomic loci and subtracted it from the estimated DSB number. The overall computation can be performed by qDSB-Seq software.

## Alternative methods

iSeq provides comprehensive analysis for DSB sequencing data from raw sequencing data processing to enrichment analysis and quantification analysis. Uniquely, iSeq calculates read coverage on DSB ends by first nucleotide of reads and in strand-specific manner, these analysis cannot be done by frequently used tools, for example, SAMtools^16^ and bedtools as mentioned above. Even though there is no available software to analysis DSB sequencing data, the users may utilize the intermediate files to conduct some specific analysis. For example, the read counts can be used as input for peak finding using MACS^12^ or HOMER^17^, both of which are designed for ChIP-seq data, to identify high-frequency DSBs with peak shape (eg. enzyme-induced DSBs). The number of reads in genomic windows can be used for conducting other statistical analysis, e.g. Fisher's exact test, which can be performed in R language. Our approach is a model-free method, which gives flexibility and is able to detect DSB patterns effectively and not limited to the shape of DSB profile.

## Advantages and limitations

### Variable scale in enrichment analysis suitable for different DSB sources

The enrichment analysis in iSeq package can be performed to identify different DSB sources, which create different DSB patterns with variable scale. Restriction enzymes induce DSBs on their recognition sequences, where their length varies from 4 bp to 50 bp based on The Restriction Enzyme Database^18^. However, DNA replication-related DSBs may be located on a wider area from a couple of kilo base pairs^13^ to mega base pairs^4^. Because enrichment analysis in iSeq is on the basis of statistical test, it is not limited by the shape of DSB patterns. Small scale can be used to detect high frequency DSBs, such as at enzyme recognition sites. On the contrary, large scale can detect broad DSBs with low DSB frequency, such as some spontaneous DSBs generated from aberrant replication or transcription^19^. Using enrichment analysis in iSeq package, we identified off-target sites for I-SceI enzyme digestion in an engineered yeast strain (YBP-275)^5^, it shows the ability of iSeq in identifying off-target sites of restriction enzymes.

### Evenly distributed DSBs are not suitable for enrichment analysis

The function of enrichment analysis can be used to identify enriched regions with significantly-increased DSBs. However, when DSBs are evenly distributed, the hypergeometric test used in enrichment analysis is not capable of identifying these DSBs. For example, we characterized Zeocin-induced DSBs by DSB sequencing^13^, even though based on our quantification results, Zeocin-induced DSBs distributed along the whole genome, we did not observe significant difference between Zeocin and control data when the sequencing reads were normalized to the total number of reads or used for enrichment analysis. Therefore, DSB quantification is required in the samples with evenly distributed DSBs.

**Advantages:**

1. **Dedicated approach for DSB detection and analysis**: adapted to unique properties of such data such as low signal, possible aggregation of reads in one place, lack of specific peak shape (depends on presence of DNA repair by Homologous Recombination and time point).
2. **Multiscale analysis:** allows the user to select the resolution that best suits the problem and the data, e.g. cytoband resolution to compare DSB sequencing data with the common fragile sites observed under the microscope. It also allows the detection of not only overlaps, but also events occurring in the proximity, for example DSBs near, but not at, stalled forks or R-loops. Multiscale analysis also eliminates spurious correlations, present only at a single resolution.
3. **Built-in data reduction scheme**: data is automatically binned and presented on a simplified grid, the same way for different data sets, facilitating comparisons.
4. **Empirical probabilities:** more reliable than theoretical, since genomes are too complex to describe mathematically.
5. **Comprehensive normalization:** Performing such stringent corrections produces different and more biologically realistic results.

### Level of expertise required

The software has been made user-friendly and has been equipped with an extensive manual and command-line help. We have provided supporting webpages for iSeq (http://breakome.utmb.edu/software.html), qDSB-Seq (https://github.com/rowickalab/qDSB-Seq), and GREDSTAT (http://gredstat.rowickalab.org). Performing the step-by-step protocol requires the basic skills in manipulating Linux that can be learned within one hour for the basic commends.

## Materials

**Equipment:**

- This package is supported for Linux operating systems. The package has been tested on the following systems:

~~~
Linux: Fedora 20
~~~
- The iSeq package requires a computer with enough RAM to support the operations defined by a user. The RAM depends on how big the datasets and the genome size are. For optimal performance, we recommend a server with the following specs:

~~~
RAM: 128+ GB
CPU: 8+ cores, 3.1+ GHz/core
HD: 100 GB of free space
~~~

**Software:**

- C++(gcc) compiler: The latest version of C++(gcc) compiler should be installed (generally comes with Linux operating system, just needs to be updated), C++ libraries called boost (www.boost.org) and gsl (www.gnu.org/software/gsl) have to be installed as well.
- PERL: 5.18.4 or higher, usually it is already installed on Linux, if not, please download and install it from https://www.perl.org/.
- R: version 3.5.1 or higher required, to install R:

1. Download R from http://cran.us.r-project.org/, click “Download R for Linux” to download the latest version.
2. Install R. Leave all default settings in the installation options.
- R package dependencies: once R is installed, type ‘R’ to enter into console, install the packages needed:

1. install.packages(“optparse”, “GenomicRanges”, “regioneR”, “ggsci”)
2. The versions of packages are: optparse: 1.6.0, GenomicRanges: 1.34.0, regioneR: 1.14.0
- Python: version 2.7 or higher with bitarray 0.8.1.
- MatLab: R2012a or higher.
- Bowtie: 0.12.2 or higher (http://bowtie-bio.sourceforge.net/index.shtml).
- FastQC: lastest (https://www.bioinformatics.babraham.ac.uk/projects/fastqc/).
- iSeq package: please visit http://breakome.utmb.edu/software.html to download *hygestat_bless*, *hygestat_annotation*, *hygestat_genes*, *hygestat_plots*, *hygestat_windows, hygestat_mappability* and install them. For *hygestat_bless*, install it by:

~~~
./configure --prefix=/path/to/install/ --with-bowtie=/path/to/bowtie/ --with-
fastqc=/path/to/fastqc/
make
make install
~~~ For *hygestat_genes* and *hygestat_windows*,

~~~
make
cp *hygestat_genes hygestat_windows* /path/to/install
~~~ For *hygestat_annotation*,

~~~
cp hygestat_annotation.1.0.py /path/to/install/
~~~
- qDSB-Seq code: visit https://github.com/rowickalab/qDSB-Seq for the information.

To install the package, use git clone:

~~~
git clone https://github.com/rowickalab/qDSB-Seq.git
~~~
or download the package and then unzip:

~~~
unzip qDSB-Seq-master.zip
~~~
Compile btt software that converts bowtie output (gcc required):

~~~
cd src
make
~~~
Export the directory of qDSB-Seq to environment:

~~~
export qDSB-Seq_DIR=ABSOLUTE_PATH_OF_QDSB_SEQ
~~~

**Input data files:**

- Genome sequence file for yeast: obtained from UCSC Genome Browser Gateway (http://hgdownload.cse.ucsc.edu/goldenPath/hg19/bigZips/chromFa.tar.gz).
- Genome sequence file for yeast: obtained from UCSC Genome Browser Gateway (http://hgdownload.soe.ucsc.edu/goldenPath/sacCer3/bigZips/chromFa.tar.gz).
- Bowtie index of *Homo sapiens* genome: the bowtie index of human genome (GRCh37) was built by the command:

~~~
bowtie-build hg19.fasta hg19.bowtie
~~~
- Bowtie index of *Saccharomyces cerevisiae* genome: the bowtie index of yeast genome (sacCer3) was built by the command:

~~~
bowtie-build sc3.fasta sc3.bowtie
~~~
- DSB sequencing data for human: we used DSB sequencing data obtained using BLESS from aphidicolin-treated (treatment) and untreated (control) U2OS-DRH-1 cells for the analysis. The aphidicolin-treated BLESS data can be downloaded from NCBI SRA database under the accession numbers: SRR695402 and SRR695426. They were merged into a fastq file, A.fastq. The control data can be downloaded under the accession numbers: SRR695321 and X. They were merged into C.fastq.
- DSB sequencing data for yeast: we used i-BLESS data from hydroxyurea (HU)-treated and untreated samples. They can be downloaded from NCBI SRA database under the accession numbers: SRR8786919 and SRR8786918, respectively.
- gDNA sequencing data for yeast: we used two datasets for HU-treated and untreated samples. They are whole genome sequencing data used for estimating cutting efficiency and DSBs per cell. They can be downloaded from NCBI SRA database under the accession numbers: SRR8786925 and SRR8786924, respectively.
- Genome mappability files: we used the genome mappability files to exclude unmappable regions from the analysis. These files were made by mapping artificial sequencing reads, which were generated from reference genome sequence using a 45 bp sliding window stepping by 1 bp, to the reference genome using bowtie with parameters ‘--concise -l45 -n0 -a -m2’. Because *hygestat_mappability* could spend a couple of hours to build mappability files, we provide pre-built mappability files for human (GRCh37), mouse (GRCm38), and budding yeast (sacCer3).
- Fragile bands and cyto bands: The human fragile bands and cyto bands can be downloaded from http://breakome.utmb.edu/software.html by clicking supplementary_files.tar.gz.

**Annotation files:**

- Genomic features: We are interested in DSB-enriched regions not only their positions but also their annotations. We use *hygestat_annotation* software to annotate DSB-enriched regions identified by *hygestat_bless* to the known genomic features. The genomic features should be BED format, including columns: chromosome, start position, and end position.
- Enzyme cutting site files: we used restriction enzyme NotI to create spike-ins in yeast cells. The cutting sites can be obtained from our tool, Genome-wide Restriction Enzyme Digestion STatistical Analysis Tool, GREDSTAT, at http://gredstat.rowickalab.org.
- Gene coordinates for human: obtained from GENCODE (https://www.gencodegenes.org/human/release_12.html)
- Replication origin file: the early origins with time < 25 min were used (http://cerevisiae.oridb.org/data_output.php?main=sc_ori_studies&table=Yabuki2002_ori&ext_format=&format=tab). The origin on rDNA was rejected. The origin file can be found in **Supplementary Data**.

### Procedure

#### Stage I: Starting from files required for the analysis (12 min)

1. *Create and set directories (2 min)*. Create main directory (all the analysis will be run under this directory):

~~~
mkdir work_dir
export work_dir=ABSOLUTE_PATH_OF_WORK_DIR
cd $work_dir
~~~

*Create subdirectories*:

~~~
mkdir hygestat_bless hygestat_annotation hygestat_genes DSB_quantification
~~~

2. *Create soft links for the required files or directories (10 min)*.

~~~
cd hygestat_bless
ln -s PATH/hg19.bowtie.*.ebwt ./ (Attention: bowtie index files include multiple ebwt files)
ln -s PATH/A.fastq treat.fastq
ln -s PATH/C.fastq control.fastq
ln -s MAPPABILITY_DIRECTORY mappability
ln -s FRAGILE_BAND_FILE fragile_bands.txt
ln -s CYTO_BAND_FILE cyto_bands.txt
~~~

~~~
cd $work_dir/hygestat_annotation
mkdir genomic_features; cd genomic_features
ln -s FEATURE_FILES . (Attention: FEATURE_FILES should contain chromosome, start, end in the first three columns)
~~~

~~~
cd $work_dir/hygestat_genes
ln -s MAPPABILITY_DIRECTORY mappability
ln -s GENE_GTF genes.gtf
~~~

~~~
cd $work_dir/DSB_quantification
ln -s SRR8786918_1.fastq B_WT_R1.fastq
ln -s SRR8786918_2.fastq B_WT_R2.fastq
ln -s SRR8786919_1.fastq B_WT_HU_R1.fastq
ln -s SRR8786919_2.fastq B_WT_HU_R2.fastq
ln -s SRR8786925_1.fastq G_WT_R1.fastq
ln -s SRR8786925_2.fastq G_WT_R2.fastq
ln -s SRR8786924_1.fastq G_WT_HU_R1.fastq
ln -s SRR8786924_2.fastq G_WT_HU_R2.fastq
ln -s ENZYME_CUTTING_SITE_FILE ./NotI.txt
ln -s YEAST_GENOME ./sc3.fasta
ln -s PATH/sc3.bowtie.* .
cat B_WT_R1.fastq B_WT_R2.fastq > B_WT.fastq
cat B_WT_HU_R1.fastq B_WT_HU_R2.fastq > B_WT_HU.fastq
~~~

#### Stage II: Identification of DSB-enriched regions (38 min)

This analysis extracts sequences from fastq or fasta file, trims close and distant barcodes (optional), aligns sequencing reads to the reference genome using bowtie, divides genome into intervals and calculates enrichment score using hypergeometric test, and performs multiple hypothesis testing corrections using Benjamini-Hochberg correction and Bonferroni correction.

3. *Change working directory (1 min)*.

~~~
cd $work_dir/hygestat_bless
~~~

4. *Run hygestat (26 min)* to do hypergeometric test by sliding window. Create a bash file *work.sh* with content:

~~~
hygestat -o fastq \
-O hygestat_treat_vs_control \
-p 5 \
-G hg19.bowtie \
-g human \
-F control.fastq -N bless \
-f treat.fastq -n bless \
-m mappability/ \
-k -r 48000 -w 1 \
-y cyto_bands.txt \
-Y fragile_bands.txt > hygestat_bless.log 2>&1
~~~

Then, run the command:

~~~
sh work.sh
~~~

5. *Count significant intervals under aphidicolin treatment. (2 min)*

5.1 *Count significant intervals based on Q-value corrected by Benjamin-Hochberg method*.

~~~
awk ‘$7<0.05’ treat_vs_control_close_barcode/48000_hygestat_treat_vs_control | wc-l
~~~

5.2 *Count significant intervals based on Q-value corrected by Bonferroni method*.

~~~
awk ‘$8<0.05’ treat_vs_control_close_barcode /48000_hygestat_ treat_vs_control |wc -l
~~~

6. *Plot chromosome ideogram and significant intervals using hygestat_plots (6 min)*.

6.1 *Enter into the directory of enrichment analysis and copy files.*

~~~
cd treat_vs_control_close_barcode
cp produce_chromosome_plot.m chromosomeplot2.m hs_cytoBand_hg19.txt ./
~~~

6.2 *Run generate_data_files.sh in hygestat_plots to generate interval files by chromosome (1 min)*

~~~
sh generate_data_files.sh 48000 48000_hygestat_treat_vs_control
~~~

6.3 *Open MATLAB console and click “Open” and choose produce_chromosome_plot.m script file, then click “Run” (5 min).*

#### Stage III: Annotation of DSB-enriched regions (2 min)

This analysis computes the feature score and enrichment *P*-value by permutation test to evaluate how DSB regions are enriched in the given features.

7. *Change working directory (1 min)*.

~~~
cd $work_dir/hygestat_annotation
~~~

8. *Run hygestat_annotation (1 min)*. When the mappability data is provided, the analysis is performed only in the mappable genome regions, which is recommended for the sequencing data to increase accuracy. Here, we use annotation of DSB-enriched regions to CpG islands as an example:

~~~
hygestat_annotation.1.0.py -F genomic_features/CpGislands.bed \
-B ../hygestat_bless/treat_vs_control_close_barcode/48000_hygestat_treat_vs_control\
-f CpG_island \
-b treat_vs_control_48000 -O treat_vs_control_48000.annotations \
-p 0.05 -c 8 -n 1000
~~~

#### Stage IV: (optional) Identification of DSB-enriched genes (2 min)

This analysis identifies DSB-enriched genes based on hypergeometric test. The software *hygestat_genes* performs similar computation with *hygestat_bless*. Differently, it compares treatment with control sample in gene regions rather than continuous windows.

9. *Change working directory (1 min)*:

~~~
cd $work_dir/hygestat_genes
~~~

10. *Run hygestat_genes (2 min)*.

~~~
hygestat_genes ../hygestat_bless/treat.fastq_close_barcode/ \
../hygestat_bless/control.fastq_close_barcode/ \
mappability/ genes.gtf
~~~

#### Stage V: Quantification of DSBs and 1-ended DSBs genome-wide (158 min)

We reported qDSB-Seq method to quantify DSBs by inducing spike-ins on specific loci by restriction enzyme. Here, we provide a guide how to use qDSB-Seq software to quantify DSBs per cell and DSBs genome-wide. We will use i-BLESS data from the budding yeast for an example and quantify replication-related DSBs.

11. *Change working directory (1 min).*

~~~
cd $work_dir/DSB_quantification
~~~

12. *Obtain enzyme cutting sites via GREDSTAT* (2 min). Visit http://gredstat.rowickalab.org, click Virtual digestion, choose organisms and enzymes and run. Download enzyme cutting sites. We also provide NotI cutting sites in **Supplementary Data**. (Attention: The coordinates of cutting sites should be downstream of cutting sites on Watson and Crick strands.)

13. *Generate random genome coordinates (1 min)*. We used random genome coordinates to deplete background noise in cutting efficiency calculation. First, length of each chromosome should be obtained:

~~~
perl $qDSB_Seq_DIR/PERL/fastaLength.pl sc3.fasta > sc3.length
~~~

Then, select genomic coordinates randomly from command line for later use to subtract background from cutting efficiency:

~~~
Rscript $qDSB_Seq_DIR/R/generate_background_bed.R \
sc3.length 1000 background.bed
~~~

14. *Run hygestat in Step 4 to generate read mapping output (30 min)*.

~~~
hygestat -o fastq \
-O hygestat_WT_HU_vs_WT \
-p 5 \
-G sc3.bowtie \
-g yeast \
-F B_WT.fastq -N bless \
-f B_WT_HU.fastq -n bless \
-m mappability/ \
-k -r 1000 -w 1 > hygestat_bless.log 2<&1
This software will produce two sequence files we need for the further analysis:
B_WT_HU.fastq.dump.fa.TCGAGGTAGTA.fasta.bt.btt and
B_WT.fastq.dump.fa.TCGAGGTAGTA.fasta.bt.btt
~~~

15. *Prepare gDNA sequencing files (12 min)*. Our gDNA data for WT_HU sample (SRR8786924) includes multiple sequencing libraries, then it is huge. Here, we only take the first 39734298 reads for the analysis.

~~~
head -158937192 G_WT_HU_R1.fastq > G_WT_HU_R1.158937192.fastq
head -158937192 G_WT_HU_R2.fastq > G_WT_HU_R2.158937192.fastq
~~~

Convert fastq files into sequence files:

~~~
perl $qDSB_Seq_DIR/PERL/fastq2seq.pl G_WT_HU_R1.158937192.fastq >
G_WT_HU_R1.seq
perl $qDSB_Seq_DIR/PERL/fastq2seq.pl G_WT_HU_R2.158937192.fastq >
G_WT_HU_R2.seq
perl $qDSB_Seq_DIR/PERL/fastq2seq.pl G_WT_R1.fastq > G_WT_R1.seq
perl $qDSB_Seq_DIR/PERL/fastq2seq.pl G_WT_R2.fastq > G_WT_R2.seq
~~~

16. *Run qDSB-seq.pl (89 min)* to quantify total DSBs per cell and DSBs per million cells on each genomic position.

~~~
perl $qDSB_Seq_DIR/qDSB-seq.pl \
B_WT_HU.fastq.dump.fa.TCGAGGTAGTA.fasta \
G_WT_HU_R1.seq G_WT_HU_R2.seq -s WT_HU -r ‘WT HU’ -g yeast \
-f sc3.fasta -i sc3.bowtie -e NotI -t 5 -c NotI.bed -b background.bed -p WT_HU \
-h B_WT_HU.fastq.dump.fa.TCGAGGTAGTA.fasta.bt.btt
perl $qDSB_Seq_DIR/qDSB-seq.pl \
B_WT.fastq.dump.fa.TCGAGGTAGTA.fasta G_WT_R1.seq G_WT_R2.seq \
-s WT -r ‘WT’ -g yeast -f sc3.fasta -i sc3.bowtie -e NotI -t 5 -c NotI.bed \
-b background.bed -p WT \
-h B_WT.fastq.dump.fa.TCGAGGTAGTA.fasta.bt.btt
~~~

(Attention: The parameter “-t” is the DSB end type created by restriction enzyme. The default is 3’-overhang, set it to “-t 5” if the enzyme creates 5’-overhang and use default for blunt ends.)

17. *Draw barplot for the across-samples comparison of the total number of DSBs (2 min).*

17.1 *Prepare data for barplot (1 min)*.

~~~
cat WT.DSBs.summary.txt > DSBs.summary.txt
cat WT_HU.DSBs.summary.txt | grep –v sample_name >> DSBs.summary.txt
~~~

17.2 *Plot DSBs (1 min)*.

~~~
Rscript $qDSB_Seq_DIR/R/plot_DSBs.R DSBs.summary.txt
~~~

18. *Draw DSB profile around replication origins (14 min)*.

18.1 *Generate data matrix around replication origins (12 min)*.

~~~
Rscript $qDSB_Seq_DIR/R/prepare_data_metaprofile.R WT.DSBs_perM.bedGraph \
origins.bed WT.metaprofile 10000 10000 1000 1000
~~~

~~~
Rscript $qDSB_Seq_DIR/R/prepare_data_metaprofile.R \
WT_HU.DSBs_perM.bedGraph origins.bed WT_HU.metaprofile \
10000 10000 1000 1000
~~~

18.2 *Build data for plot (1 min)*.

~~~
Rscript $qDSB_Seq_DIR/R/build_data.R WT.metaprofile.mat WT.metaprofile \
WT WT WT
Rscript $qDSB_Seq_DIR/R/build_data.R WT_HU.metaprofile.mat \
WT_HU.metaprofile WT_HU WT_HU WT_HU
~~~

~~~
Merge results from WT and WT HU:
cat WT.metaprofile.txt WT_HU.metaprofile.txt > metaprofile_WT_WT_HU.txt
~~~

18.3 *Plot metaprofile around replication origins (1 min)*.

~~~
Rscript $qDSB_Seq_DIR/R/plot_lines.R \
metaprofile_WT_WT_HU.txt metaprofile_WT_WT_HU.txt "Distance to origin" \
"DSBs per mln cells" -10000 10000 0 50
~~~

19. *Quantification of 1-ended DSBs (7 min)*.

This analysis identifies and quantifies 1-ended DSBs using hypergeometric test on Watson and Crick strands.

19.1 *Prepare read counts on plus (Watson) and minus (Crick) strands (2 min).*

~~~
perl $qDSB_Seq_DIR/PERL/createChrFiles.pl \
B_WT.fastq_close_barcode/ B_WT.fastq_distant_barcode/ B_WT
~~~

~~~
perl $qDSB_Seq_DIR/PERL/createChrFiles.pl \
B_WT_HU.fastq_close_barcode/ B_WT_HU.fastq_distant_barcode/ B_WT_HU
~~~

19.2 *Run hypergeometric test for Watson and Crick strands (2 min)* to identify windows with significant difference between both strands.

~~~
mkdir B_WT.plus_vs_minus B_WT.minus_vs_plus \
B_WT_HU.plus_vs_minus B_WT_HU.minus_vs_plus
~~~

Run hygestat_windows compiled from *hygestat_windows*:

~~~
hygestat_windows yeast \
B_WT.plus_strand/close_barcode/ \
B_WT.minus_strand/close_barcode/ \
B_WT.plus_vs_minus/output_500.txt 500 501
~~~

~~~
hygestat_windows yeast \
B_WT.minus_strand/close_barcode/ \
B_WT.plus_strand/close_barcode/ \
B_WT.minus_vs_plus/output_500.txt 500 501
~~~

~~~
hygestat_windows yeast \
B_WT_HU.plus_strand/close_barcode/ \
B_WT_HU.minus_strand/close_barcode/ \
B_WT_HU.plus_vs_minus/output_500.txt 500 501
~~~

~~~
hygestat_windows yeast \
B_WT_HU.minus_strand/close_barcode/ \
B_WT_HU.plus_strand/close_barcode/ \
B_WT_HU.minus_vs_plus/output_500.txt 500 501
~~~

19.3 *Obtain read counts from predicted 1-ended DSBs (1 min)*.

~~~
cat B_WT.plus_vs_minus/output_500.txt \
B_WT.minus_vs_plus/output_500.txt \
| perl $qDSB_Seq_DIR/PERL/generate_bedGraph_for_1DSB.pl 0.05 \
BF B_WT
~~~

~~~
cat B_WT_HU.plus_vs_minus/output_500.txt \
B_WT_HU.minus_vs_plus/output_500.txt \
| perl $qDSB_Seq_DIR/PERL/generate_bedGraph_for_1DSB.pl 0.05 \
BF B_WT_HU
~~~

19.4 *Quantify 1-ended DSBs (1 min)*.

~~~
Rscript $qDSB_Seq_DIR/qDSB-seq.R -s WT -e NotI -g WT \
-m process_gDNA_data/cutting_efficiency_NotI/WT.NotI.L0-R0.exact.switch.txt \
-b
~~~

~~~
process_gDNA_data/cutting_efficiency_NotI/WT.NotI.background.exact.switch.txt \
-r process_DSB-seq_data/get_NotI_reads/WT.NotI.flank_0_0.txt \
-d B_WT.1DSB.bedGraph -t 1 -c 10 -n 0 -a 1 -p WT.1DSB
~~~

~~~
Rscript $qDSB_Seq_DIR/qDSB-seq.R -s WT_HU -e NotI -g WT_HU \
-m process_gDNA_data/cutting_efficiency_NotI/WT_HU.NotI.L0-
R0.exact.switch.txt \
-b
process_gDNA_data/cutting_efficiency_NotI/WT_HU.NotI.background.exact.switch. txt \
-r process_DSB-seq_data/get_NotI_reads/WT_HU.NotI.flank_0_0.txt \
-d B_WT_HU.1DSB.bedGraph -t 1 -c 10 -n 0 -a 1 -p WT_HU.1DSB
~~~

(Attention: The parameter “-t” is the number of DNA ends. The default is ‘-t 2’ for 2-ended DSBs. Here, it should be set to 1 for 1-ended DSBs.)

19.5 *Plot 1-ended DSBs (1 min)*.

~~~
cat WT.1DSB.DSBs.summary.txt > 1DSB.summary.txt
cat WT_HU.1DSB.DSBs.summary.txt | grep -v sample_name >> 1DSB.summary.txt
Rscript $qDSB_Seq_DIR/R/plot_DSBs.R 1DSB.summary.txt "1-DSBs per cell"
~~~

### Timing

The time estimated includes the running time of source code. It does not include the time in preparation of input files, however, it includes the time in creating directories and soft links of files before running the software. The examples were run on a Linux machine (3.1 GHz 8 cores CPU, 256 GB memory). The time highly depends on dataset used, for example, the number of sequencing reads used in Stage II, the number of features used for annotation in Stage III.

Stage I: Steps 1-2, starting from files required for the analysis: ~12 min
Stage II: Steps 3-6, identification of DSB-enriched regions: ~36 min
Stage III: Steps 7-8, annotation of enriched-DSB regions: ~6 min
Stage IV: Steps 9-10, (optional) Identification of DSB-enriched genes: ~11 min
Stage V: Steps 11-19, Quantification of DSBs and 1-ended DSBs genome-wide ~158 min

### Troubleshooting

Troubleshooting can be found in Table 1.

**Table 1.** Troubleshooting table

| Step | Problem | Possible reason | Solution |
| --- | --- | --- | --- |
| 4 | Problems in running hygestat; no output | The required dependencies of software are not installed; input files are not correctly provided | Install all C++ libraries, including boost and gsl; provide files as shown in Step 4, for example, -G is the prefix of bowtie index, -m is folder name and should be end by '/' |
| 4 | No significant intervals | The size of intervals is too small, thus the number of reads used for hypergeometric test is not enough; The significance threshold is too high | Set a bigger interval size; use Benjamin-Hochberg correction or a low significance threshold |
| 7 | The given feature is not enriched or depleted | The feature is lack of specificity; the number of loci of a feature is too few, thus it is not meaningful in statistics | Use features with specificity, for example, DNA replication-related genes rather than all genes; use the feature with more loci |
| 16 | Problems in running qDSB-seq.pl | The required R libraries are not properly installed | Make sure that all package dependencies are installed; change to the same version of R and packages |
| 16 | Extreme DSBs obtained | The cutting efficiency is not correctly estimated due to wrong parameters | Make sure correct enzyme used; Set correct value for the DSB end type created by restriction enzyme, the default "-t" is 3'-overhang |
| 19 | Unexpected 1-ended DSBs | The correction and *P*-value in identifying 1-ended DSBs determines the number of reads from 1-ended DSBs (19.3) | Change correction method and *P*-value |

### Anticipated results

If success, *hygestat* in Step 4 will output multiple files and directories, some of which are provided in **Supplementary Data**:

*\*.dump.fa*: they contains only sequences extracted from provided fastq or fasta files.
*\*.dump.fa.TCGAGACGACG(TCGAGGTAGTA).fasta*: they contains sequences with close (or distant) barcode trimmed.
*\*.dump.fa.TCGAGACGACG(TCGAGGTAGTA).fasta.bt*: they are bowtie outputs in bowtie format mapped from sequences with close (or distant) barcode trimmed.
*\*.dump.fa.TCGAGACGACG(TCGAGGTAGTA).fasta.bt.btt*: the sequences in bowtie format were converted according to 5’ -> 3’ direction.
*treat_vs_control_close(distant)_barcode*: including files of enrichment analysis, *48000_hygestat_treat_vs_control* (including 8 columns: 1^st^ chromosome name; 2^nd^ and 3^rd^ start and end coordinates of the window, respectively; 4^th^ and 5^th^ the number of reads for treated and control samples, respectively; 6^th^ *P*-value, 7^th^ Q-value based on Benjamin Hochberg correction; 8^th^ *Q*-value based on Bonferroni correction) and *treat_VS_control_at_48000.bedgraph* for visualization (including 4 columns: 1^st^ chromosome name; 2^nd^ and 3^rd^ start and end coordinates of the window, respectively; 4^th^ -log10(*P*-value). In Step 5, we identified 2345 significant intervals based on Benjamin Hochberg method, and 218 significant intervals based on Bonferroni method.
*treat.fastq_vs_control.fastq.tab*: statistical results of sequencing data.
*treat(control).fastq.csv*: statistical results of data quality.
By taking output file of *hygestat*, DSB enrichment score (−10Log_10_(q-value)) can be visualized by chromosome plot in Step 6. An example on chromosome 10 is shown in **Fig. X**.

Annotation of DSB-enriched regions in Step 8 generates feature score. The output includes 8 columns, among them *feature_score* is the proportion between observed overlap and expected overlap, when it is bigger than 1, and *pv_enrichment* is significant, it suggests DSBs are enriched in the given feature. Instead, when *feature score* is smaller than 1 and *pv_depletion* is significant, it suggests DSBs are depleted in the given feature. In our example, *feature_score* is 0.72 and *pv_depletion* is 0.04, it means DSBs are depleted in CpG island.

DSB-enriched genes were identified by *hygestat_genes* in Step 10. The resulting *output.txt* file contains 9 columns, including the number of sequencing reads in genes in treatment and control samples (columns 6^th^ and 7^th^, respectively), *P*-values calculated by hypergeometric test (column 8^th^) and *Q*-values corrected by Benjamin-Hochberg method (column 9^th^).

If success, *qDSB-seq.pl* in Step 16 will output two files, including:

*\*.DSBs.summary.txt*: it contains the total number of DSBs (column “DSBs”), the standard deviation of DSBs calculated from multiple cutting sites (column “DSBs_sd_by_site”), the standard deviation of DSBs calculated from cutting efficiency (column “DSBs_sd_by_fcut”), and the number of cutting sites used for calculation (column “nsites”).
*.DSBs_perM.bedGraph: it contains four columns chromosome, start position, end position, and DSBs per million cells in the region.

The number of DSBs estimated in WT and WT HU samples are 22 and 185, respective as shown in **Fig. 4a**. The metaprofile of DSBs around replication origins is shown in **Fig. 4b**.

**Fig. 4a.**
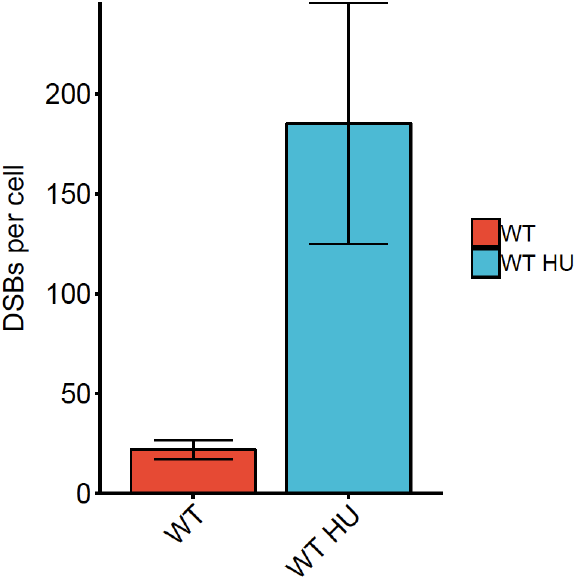
Sample plot of DSBs per cell in different samples generated in Step 17. DSBs per cell values and s.d. were calculated based on the method in qDSB-Seq paper.

**Fig. 4b.**
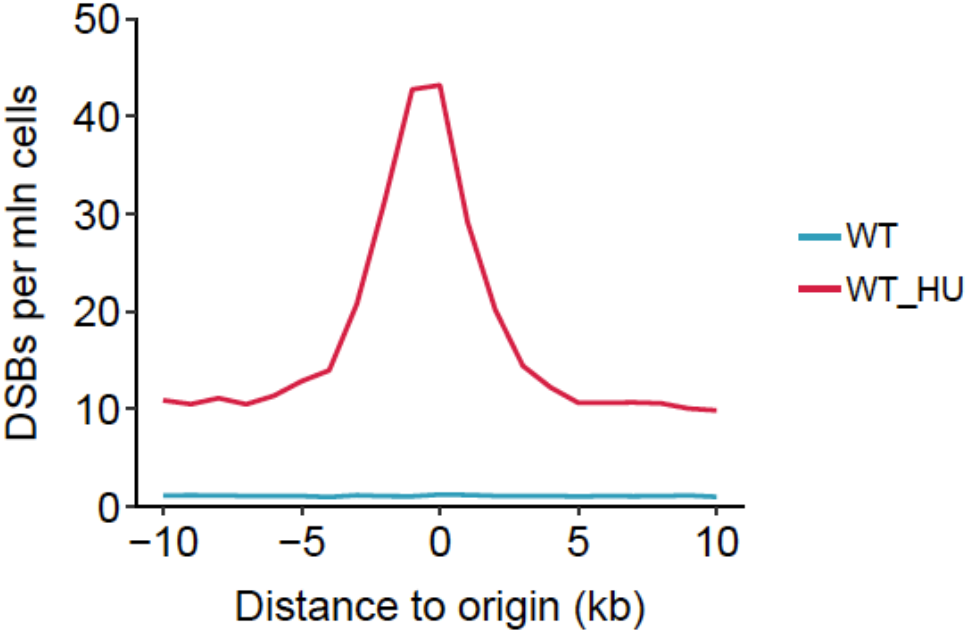
Sample plot of DSB metaprofile around origins generated in Step 18. 144 early origins were collected from Yabuki et al. Median value was taken for each bin.

We identified DSB sequencing reads from 1-ended DSBs by comparing Watson and Crick strand reads in Step 19. If success, Step 19.3 will generate a *bedGraph* file end by *1DSB.bedGraph*, it contains reads from 1-ended DSBs. Using this file, the number of 1-ended DSBs will be calculated by *qDSB-seq.R* in Step 19.4. The results in our examples are shown in **Fig. 5**.

**Fig. 5.**
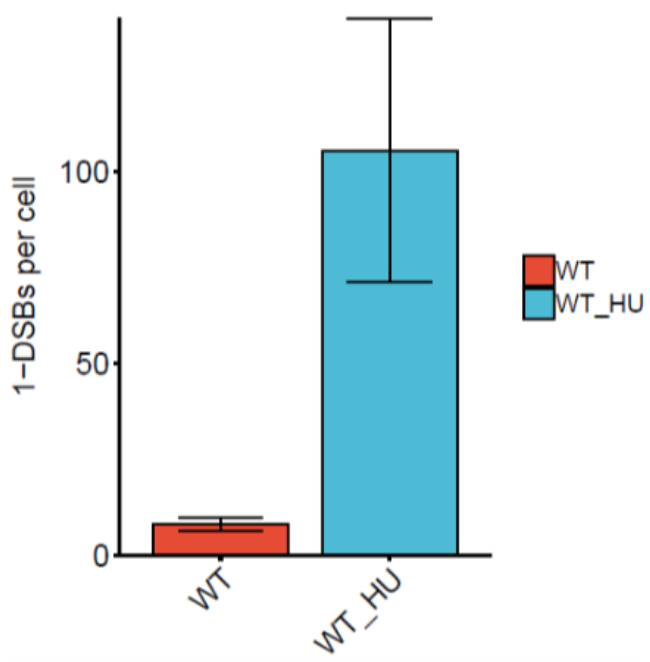
Sample plot of 1-DSBs per cell in different samples generated in Step 19. 1-DSBs per cell values and s.d. were calculated based on the method in qDSB-Seq paper

## Author contributions

M.R. supervised and coordinated the project. A.B. initially developed core algorithms in iSeq package. N.D. developed the method for annotation analysis. B.F. and J.N. integrated iSeq package. Y.Z. developed the qDSB-Seq software for quantification analysis. Y.Z. and M.R. wrote the manuscript. Y.Z. performed data analysis and prepared figures. R.Y. contributed to software development and data analysis. All authors read the manuscript.

## Acknowledgements

This research was supported by the NIH grant R01GM112131 to M.R. (Y.Z., N.D., B.F., J.N., R.Y. and M.R.). This work was also supported by a training fellowship from the Gulf Coast Consortia on the Computational Cancer Biology Training Program (CPRIT Grant No. RP170593) to Y.Z, and National Science Center grant 2016/21/B/ST6/01471 to N.D.

---

**Supplementary Figure 1.**
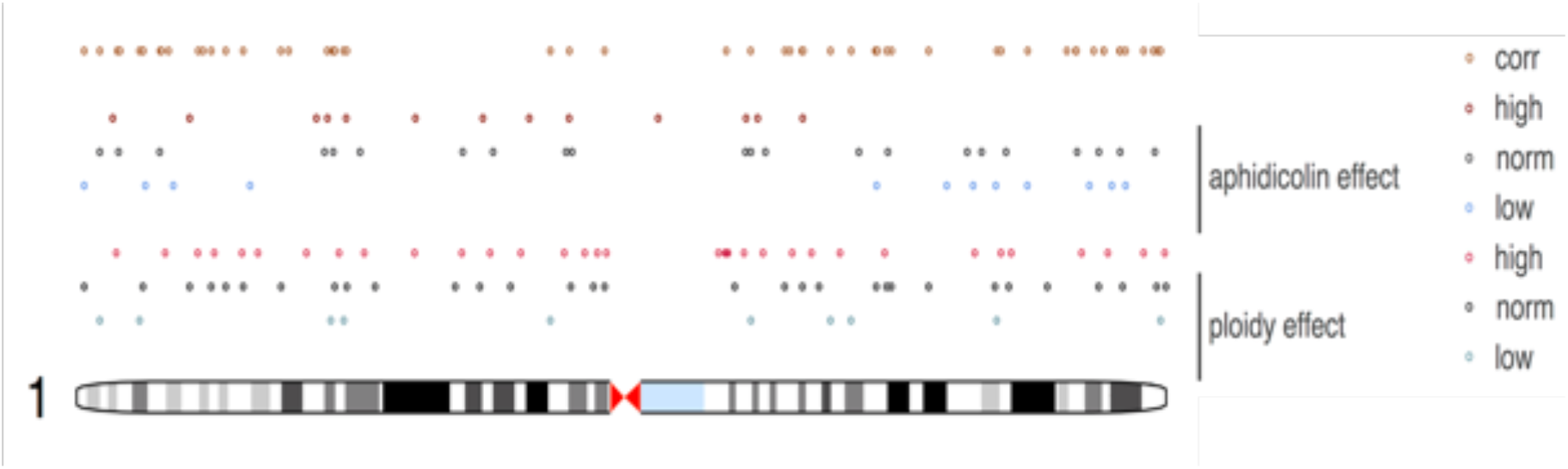
Effects of CNV (ploidy), treatment (aphidicolin) and normalized DSB-prone regions for human chromosome 1. Note that due to the comprehensive corrections, DSB-prone regions (corrected, top row) do not coincide with genomic regions over-represented in sequencing due changes in replication induced by aphidicolin treatment (2^nd^ row from the top) or due to CNV in HeLa cells (5^th^ row from the top).

**Supplementary Figure 2.**
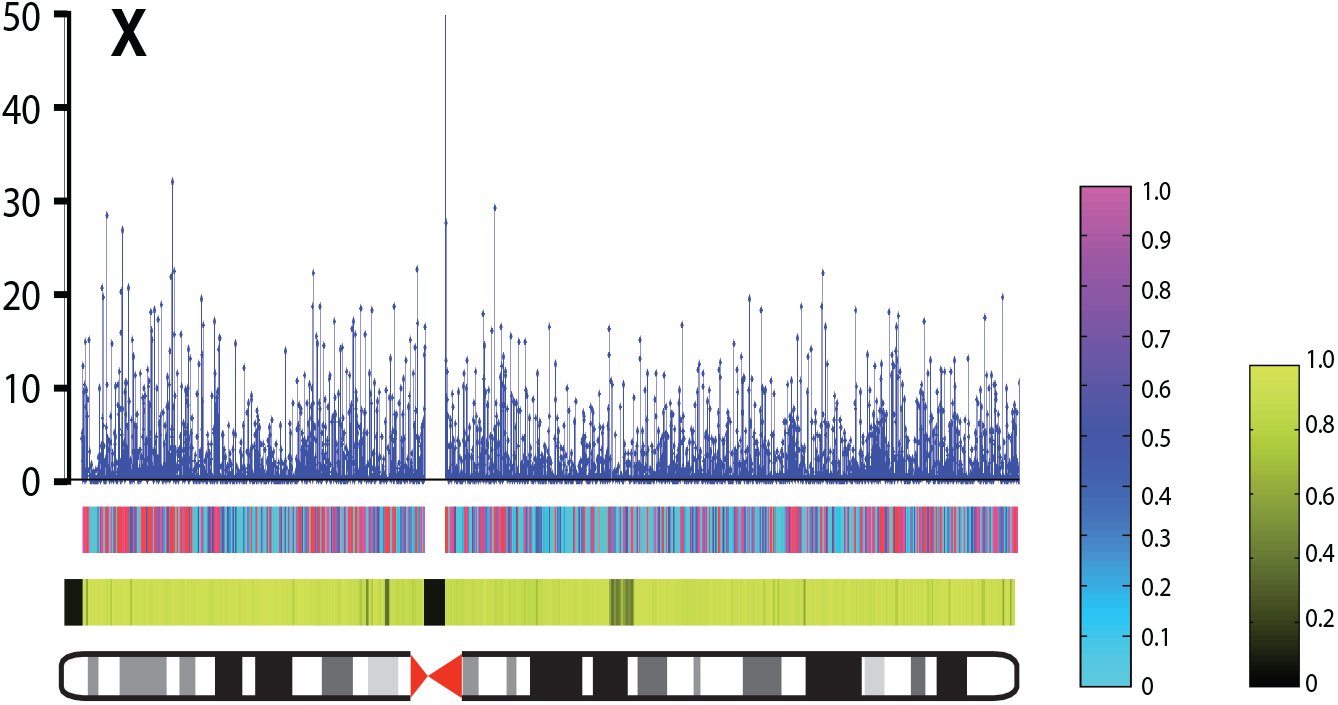
Example of multiscale fragility maps generated by iSeq (shown aphidicolin-induced DSBs in HeLa cells, chromosome X). From bottom: 1) chromosome ideogram (centromere is denoted by red hourglass); 2) mapability (black denotes 0%); 3) breaking probability (purple denotes 100%, light blue 0%) for 200 kilobase resolution; 4) −10Log_10_(*Q* values) for DSB presence at 48 kilobase resolution (blue stem plot).

